# A preliminary assessment of the genome sequence of the Annual Mercury, *Mercurialis annua*

**DOI:** 10.1101/2024.06.24.600384

**Authors:** Mratunjay Dwivedi, Nagarjun Vijay

## Abstract

**Summary:** Review of Christenhusz et al., Wellcome Open Research. A high-quality genome sequence of the Annual Mercury, *Mercurialis annua* is generated as part of the Darwin Tree of Life Project. The Annual Mercury plant is a good study system for evolutionary transitions between sexual systems, mechanisms of sex determination in plants and changes in ploidy level. The XX female sequenced in this study provides a chromosome-level assembly with reasonable contiguity and high gene completeness. We compare the gene completeness of the assembly presented with those of closely related species, including another published assembly of the Annual Mercury generated using a polyploid individual. Annotation and comparative analysis of the organelle genomes suggest complete circular assemblies. However, we note that the NCBI submissions are tagged as linear. The repeat content identified on the autosomes and X chromosome appears comparable, suggesting the sex chromosomes are relatively recent. An alternative possibility is that our preliminary analysis failed to identify repeats unique to the X chromosome or that these repeat regions have not been assembled in the genome. The harder-to-assemble centromere and telomere regions are not annotated in the genome and are potentially incomplete or missing. We identify putative centromere regions with elevated GC content, but they must be validated. The demographic histories reconstructed for the autosomes and X suggest distinct trajectories irrespective of the scaling parameters. Compared to the autosomes, the older histories recorded on the X are a promising avenue for further work to study the origin of the sex chromosome. Our preliminary assessment of the genome sequence suggests the genome is of sufficiently good quality for use as a reference for diverse analysis aimed at answering important eco-evolutionary questions of interest that can be answered using this system.

## Introduction

Christenhusz et al. present a chromosome-level genome assembly from a diploid female *Mercurialis annua* (Annual Mercury) obtained by assembly of short and long-read sequencing data as part of the Darwin Tree of Life Project (Blaxter 2022). The importance of Annual Mercury as a model for studying evolutionary transitions between sexual systems (Durand 1963), mechanisms of sex determination in plants (Westergaard 1958) and ploidy levels (Russell and Pannell 2014) provides the necessary rationale for generating the genome assembly. Christenhusz et al. provide a comprehensive list of eco-evolutionary questions of interest that can be answered using this system. For instance, it is mentioned that Professor. John R. Pannell became interested in this species to study “the ecological and genetic reasons for the evolution and maintenance of dioecy.”(Sánchez Vilas and Pannell 2011; Gerchen et al. 2022).

The contrast in phenotypes (presence/absence of a perennating rhizome, differences in branching frequency and colour, ecological preferences for more/less shady sites) between *M. annua* and its congener *Mercurialis perennis* (Dog’s Mercury) provides a well-suited system for comparative genomics. While an assembly for *M. perennis* is currently unavailable in the public databases, sequencing of its genome would be an excellent subsequent target for comparative genomics. Although ten accepted species are listed in the genus *Mercurialis, M. perennis* and *M. annua* are the most well-studied (Mukerji 1936). Notably, accidental plant poisoning due to *M. perennis* is reported in the literature, which ends with a cautionary note stating, “Publications which fail to give accurate and detailed colour illustrations of plants should be treated with caution.”(Rugman et al. 1983). It would be worth exploring whether *M. annua* is also poisonous.

Christenhusz et al. provide a colour photograph (**Figure 1**) of male and female *M. annua* for accurate species identification. It would be good to update the figure legend to clarify that the male plant is on the right side and the female plant is on the left or separate the photographs, as shown by Khadka et al. (Khadka et al. 2019). The genome assembly generated in this study is from an XX female. The plans to sequence a genome from a male (to study differences between the X and Y chromosomes) explain how this genome fits into the broader scheme of things. In this open review of the Christenhusz et al. manuscript, we compare the quality of the newly generated genome assembly with those available for the plants from the Acalypheae tribe, annotate the organelle genomes and compare them with those of related species, quantify and compare repeat abundance between the autosome and X, predict putative centromeres and infer demographic history using PSMC. During our preliminary analysis, we identified some minor issues related to metadata documentation that the authors of the primary study can resolve.

**Fig. 1.**
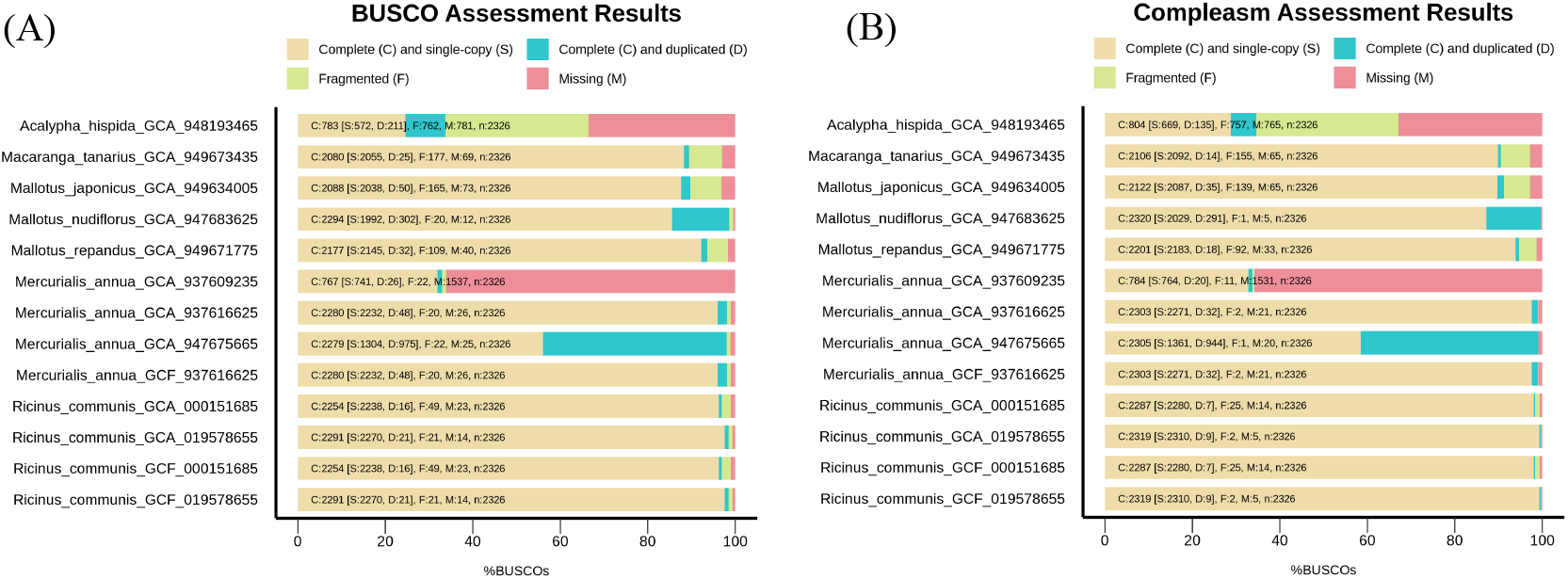
Comparative assessment of gene completeness among Acalypheae genomes using Eudicotyledons_odb10 dataset. **(A)** BUSCO-based assessment **(B)** Compleasm-based assessment. *Mercurialis annua* genome has a relatively complete assembly compared to other species.

## Methods

### Dataset and meta information

The raw sequencing datasets are provided as part of the BioPorject # PRJEB50969 and consist of Hi-C, RNA-Seq and WGS (Sequel II and Illumina NovaSeq 6000). The Illumina data comprises two BioSamples (SAMEA7522241 and SAMEA7522246) with identical meta information. It is unclear whether these BioSamples refer to two separate plants or how they differ. This information will be crucial for subsequent analysis of this “Open Data”. Another BioSample ID (SAMEA7522176) is used in Table 1 and is linked to all the sequencing runs in the metainformation, which adds to the confusion regarding the BioSample information.

### Comparative assessment of Acalypheae genome quality

We downloaded all the genome assemblies available on NCBI for the Acalypheae tribe from the subfamily Acalyphoideae under the family Euphorbiaceae. The currently available genomes include one chromosomal level(Lu et al. 2022) and several draft genome assemblies of the castor bean assembly (*Ricinus communis*). During the review, it was noticed that another genome assembly was reported for *M. annua* by Long et.al., (Long et al. 2023). The same Long et.al., (Long et al. 2023) study generates draft genome assemblies for several other Euphorbiaceae species, with five species (*Mallotus japonicus, Mallotus nudiflorus, Mallotus repandus, Macaranga tanarius* and Acalypha hispida) from the Acalypheae tribe. To asses the completeness of the genomes in terms of gene completeness, we used the BUSCO (Simão et al. 2015) tool to estimate the fraction of Complete (single copy & duplicated), Fragmented and Missing genes using the eudicots_odb10 dataset. The recent availability of compleasm (Huang and Li 2023), a faster and more comprehensive tool to assess gene content in genome assemblies, has motivated reviewers (Hiller 2024) to suggest using this tool. We use both BUSCO and compleasm on this set of 13 genomes to obtain a comparartive assessment of genome qualities to better understand the quality of the *M. annua* genome assembly generated by Christenhusz et al.

### Organelle genome annotation

It is noted that the “mitochondrial genome is 435.28 kilobases in length, while the plastid genome is 169.65 kilobases in length”. However, no attempt has been made to annotate these organelle genomes. As a quality control measure, we argue that annotating these genomes should be an essential requirement for submitting these genomes to NCBI. The methods note that “A representative circular sequence was selected for each from the graph based on read coverage.”. However, the NCBI GenBank file is tagged as a linear DNA for both the mitochondrion and plastid:chloroplast. We annotate the mitochondrion and chloroplast genomes using GeSeq and visualise them using OGDRAW implemented in CHLOROBOX(Tillich et al. 2017; Greiner et al. 2019). Detailed settings used and outputs are documented in the GitHub repository.

### Estimation of repeat abundance

The repeat regions in the genome were identified using RepeatMasker (Smit, AFA, Hubley 2015) to annotate repeat types, tabulate their abundance and mask the repeat sequences. The Kimura two-parameter divergence estimates were calculated based on the RepeatMasker.align output using accessory scripts (buildSummary.pl, calcDivergenceFromAlign.pl, and createRepeatLandscape.pl). These data were summarised as histograms depicting the distribution of repeat families. Although the global repeat landscapes seem similar between the autosomes and X, theory predicts that heteromorphic sex chromosomes will accumulate repeats, which has been demonstrated empirically (Bellott et al. 2010). To evaluate this further, we quantified the fraction of repeat content in 50Kb, 10Kb, 1Kb, 500 bp, and 100 bp windows along the length of the genome. The repeat landscapes of each chromosome were visually scanned for the presence of repeat enriched clusters.

### Demographic history inference

One of the customary genomic analyses done as soon as a genome assembly becomes available for a diploid species is the reconstruction of the demographic history of the species. In some cases, such demographic history information has been used to assess well-formulated hypotheses rigorously (Nadachowska-Brzyska et al. 2016; Gabrielli et al. 2024). Notably, genome assembly quality (Patton et al. 2019) and repeat content (Patil and Vijay 2021) have no major impact on these inferences. The short-read data associated with the *M. annua* genome were downloaded from public repositories and mapped to the genome using the bwa mem (Li 2013) read mapper. PSMC demographic inference was performed with default parameters, i.e., -t5 –r5 –p “4 + 25*2 + 4 + 6”. The default parameter settings of PSMC resulted in more than ten recombination events in each atomic interval after 20 iterations (Dataset/PSMC). Based on earlier studies, we used a mutation rate estimate of 2.5e-09 per site per year (Bai et al. 2018; Patil et al. 2021) with a generation time of 1 year to scale the PSMC inferences.

## Results

### Gene completeness in Annual Mercury is better than most other Acalypheae

GenomeScope estimated a genome size of approximately 397.24 Mb based on short-read sequencing data (Dataset/GenomeQC/genomescope). Although the genome size can vary due to the prevalence of polyploidy, the flow cytometry-based estimate reported for this individual is 690 Mb. The assembled genome is 453.2 Mb, ∼65% of the flow cytometry-based estimate and ∼114% of the GenomeScope-based estimate. The phylogenetically proximate high-quality genome of the castor bean (*Ricinus communis*) has an assembly size of 315.6 Mb. Therefore, a large fraction of the *M. annua* genome appears assembled. With a scaffold N50 of 56.1 (contig N50 of 35.6 Mb), the annual mercury genome assembly (Accession# GCA_937616625.2) by Christenhusz et al. consists of 63 scaffolds (78 contigs). Of the 63 scaffolds, eight >40 Mb scaffolds are assigned to individual linkage groups (LG-1 to LG-8). The remaining 56 scaffolds are assigned to LG4 but remain unlocalised. The reported high contiguity and gene completeness suggest a high-quality genome assembly. Our BUSCO and Compleasm-based comparison (Dataset/GenomeQC/BUSCO) with other Acalypheae genomes (**Fig. 1**) using the Eudycotyledons_odb10 dataset revealed a comparable and complete genome of *M. annua*. Notably, the earlier *M. annua* genome (Accession# GCA_947675665.1) has a high fraction of complete and duplicated BUSCOs and an assembly size of 729.6 Mb, suggesting this assembly is a polyploid. The alternate haplotype (GCA_937609235.2) assembled has a high fraction of missing BUSCOs. In summary, comparing gene completeness using BUSCO and compleasm provides highly concordant results and demonstrates the high quality of the *M. annua* genome.

### Organelle genomes seem complete and error free

The genomes of the mitochondria (OX359233.1) and chloroplast (OX359234.1) are assembled and submitted to NCBI. We provide preliminary annotation (**Fig. 2**) to identify the genes and their order in the organelle genomes (Dataset/organelle_annotation/). The *M. annua* chloroplast genome was compared one-to-one with the chloroplast genomes of *R. communis* (NC_016736.1), *A. hispida* (NC_070339.1), and *M. tanarius* (MW297079.2). The dot plot representation of these comparisons confirms a high-level sequence and gene order conservation (Dataset/dotplots). The *M. annua* mitochondrial genome could be compared only to the *R. communis* (NC_015141.1 and HQ874649.1) mitochondria, as other complete *Acalypheae* mitochondria are not currently available. Both *R. communis* mitochondria had limited similarity to the *M. annua* mitochondrial genome, mainly restricted to genic regions. The availability of additional organelle genomes from other closely related species may allow a more detailed comparison in the future. The preliminary annotations and comparisons with organelle genomes of closely related species suggest that the assembled scaffolds are complete and potentially error-free.

**Fig. 2.**
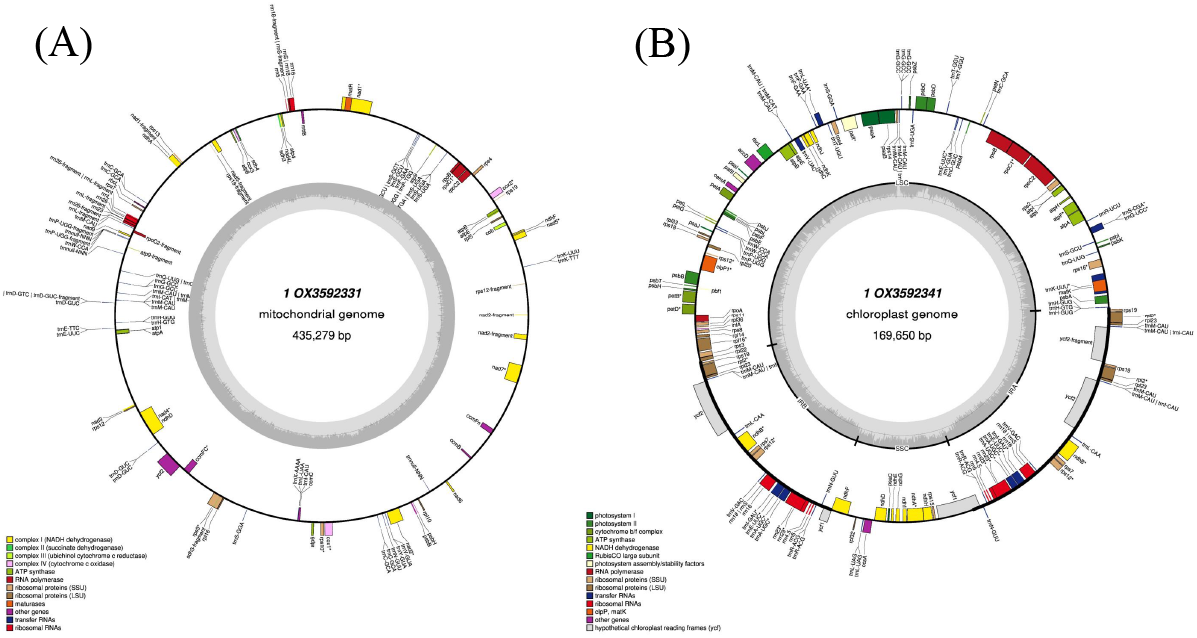
Circular plot of *Mercurialis annua* organelle genome annotation. **(A)** Mitochondria **(B)** Chloroplast. The outermost circle depicts the position and annotation of genes, and colours identify the pathway indicated at the lower left panel.

### Repeat landscapes of the autosomes and X are comparable

We compared the types of repeats, their relative abundance and prevalence in different divergence classes in the autosomes and X chromosome (**Fig. 3**). The repeat landscapes quantified based on the repeats identified by the repeat masker are highly comparable between the autosomes and X. The genomic content identified as repeat is ∼10% in both autosomes and the X. The most abundant repeat type is LTR elements (∼4%), followed by DNA transposons (< 1 %). LINE accounts for < 0.5 % of the repeat. Although sporadic windows had high repeat content, the prevalence of repeats along the length of the chromosomes did not show any prominent patterns in any of the chromosomes (Dataset/repeats/50Kb_wins). Systematic comparison of the repeat abundance in non-overlapping genomic windows of several sizes (50Kb, 10Kb, 1Kb, 500Bp, and 100Bp) failed to discern (**Fig. 4**) any differences between the autosomes and X.

**Fig. 3.**
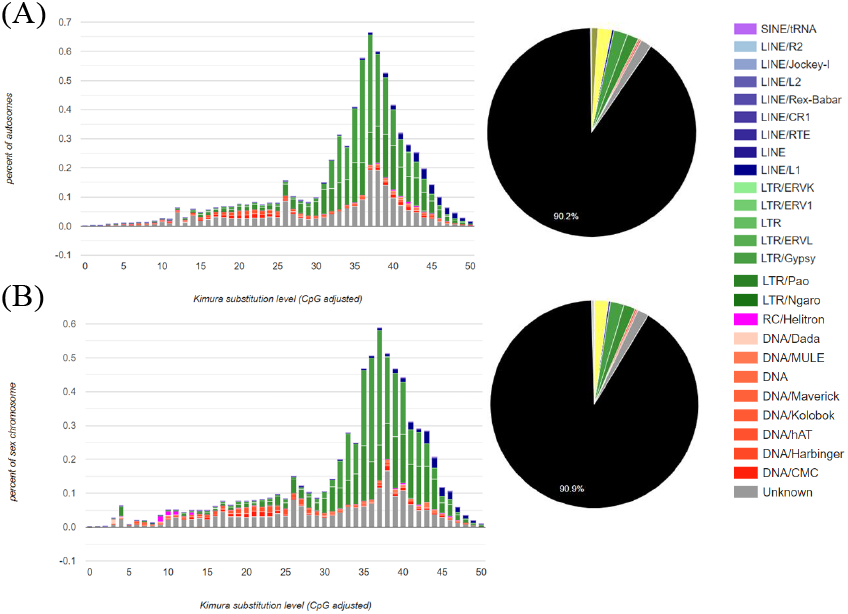
The repeat landscape of the autosomes **(A)** and the X chromosome **(B)** in the *Mercurialis annua* genome. The repeat landscapes depict the contribution of different repeat types as a fraction of the genome (along the x-axis) and their CpG-adjusted Kimura substitution level indicative of insertion age (along the x-axis). The repeats identified cover ∼10% of the assembled regions.

**Fig. 4.**
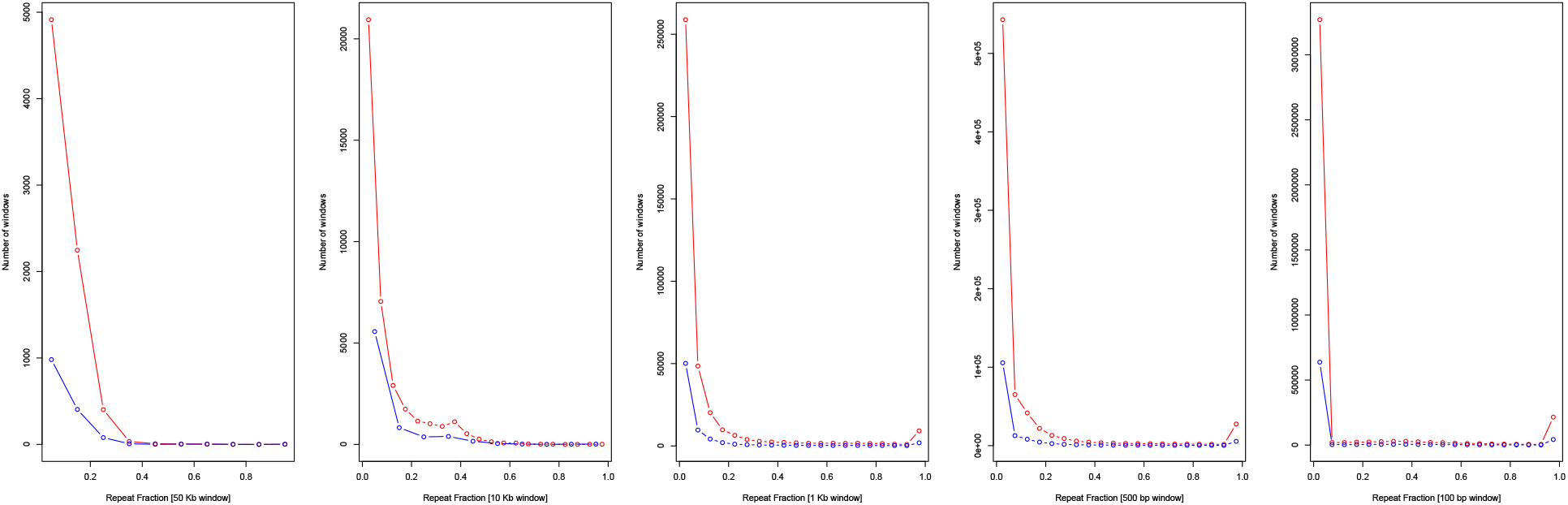
The abundance of the repeat fraction in non-overlapping genomic windows. Irrespective of the window size, the repeat fraction distributions are comparable between the autosomes (red) and X chromosome (blue).

### GC-rich candidate centromeres are assembled

Visual inspection of the GC content in non-overlapping 50Kb windows identified one prominent cluster of GC-rich windows in each chromosome except LG2 (**Fig. 5**). In LG5, in addition to the prominent cluster in the middle of the chromosome, a smaller extreme GC region occurs towards the end of the chromosome. Despite our analysis indicating this region as a putative centromere, this is not annotated in the genome assembly. We did not find any enrichment of satellite repeats near these regions, and some repeat classes were under-represented (Dataset/repeats/50Kb_wins).

**Fig. 5.**
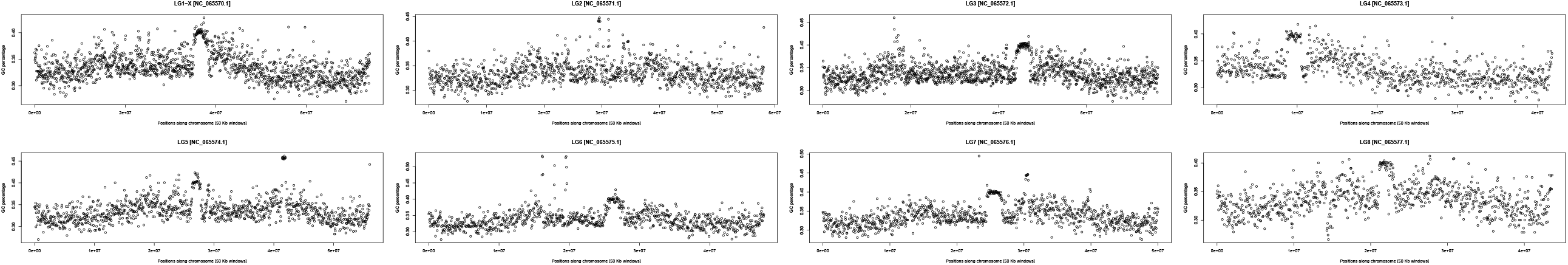
GC content variation in 50Kb non-overlapping windows along the length of each chromosome suggests putative GC-rich centromeres.

### Demographic history reconstructions suggest a recent decline in Ne

Estimation of the demographic history of the autosomes and X chromosome can provide important insights into the influence of connectivity. We use the chromosome-level genome assemblies to estimate Ne separately for the autosomes and the X (**Fig. 6**). Interestingly, the trajectory for the X chromosome extends further back in time than the autosomes, and the values of Ne are consistently higher for the X chromosome compared to the autosomes. When the results are scaled (**Fig. 6A**) with a generation time of 1 year and mutation rate of 2.5e-09 per site per year, the autosome trajectory begins around 400 Kya compared to 1200 Kya for the X chromosome. Even without scaling (**Fig. 6B**), the difference between the autosomes and X is evident.

**Fig. 6.**
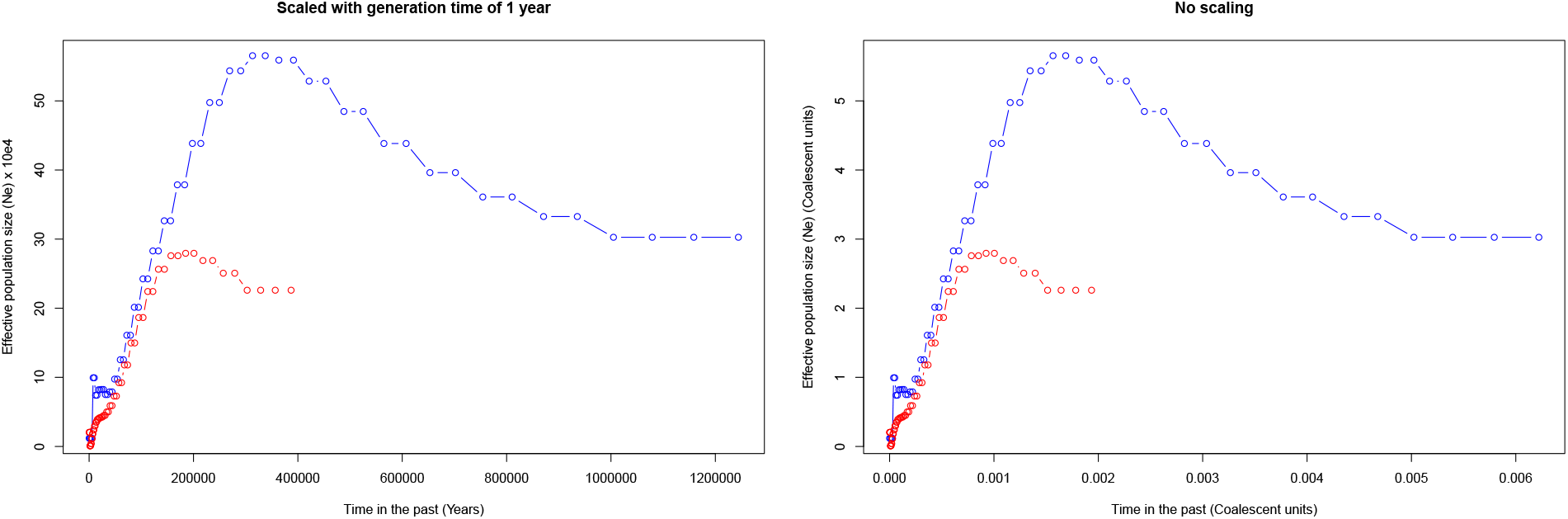
Demographic history of *Mercurialis annua* inferred using PSMC. **(A)** PSMC inferences scaled with a generation time of 1 year and mutation rate of 2.5e-09 per site per year. **(B)** PSMC inferences without any scaling (i.e., in coalescent units). The red (autosomes) and blue (X chromosome) lines represent the inferred effective population size (N_e_) at different time points.

## Discussion

In this review of the Christenhusz et al. manuscript, we include a preliminary genomic data analysis evaluating genome quality, a review of meta-data documentation, and a few subsequent genome processing steps in the spirit of Open Science. Different journals and journal-like entities have implemented some of the UNESCO recommendations on Open Science. However, research evaluation and the associated steps, such as grant funding, are still primarily a mix of the traditional impact factor-based system and the newer open science initiatives. For instance, this invitation to review provides a forum to express scientific views freely without any financial considerations. Nevertheless, publishing a data note entails article processing charges. Despite such dichotomies, Open science is helping researchers succeed (McKiernan et al. 2016).

The genome assembly quality metrics reported in the Darwin Tree of Life Project are compared to the benchmarks adapted from column VGP-2020 (Rhie et al. 2021). Our preliminary analysis compared the quality metrics with genome assemblies of other phylogenetically proximate genomes. The unique challenges of assembling diverse evolutionary lineages may not be easily amenable to benchmarks. For instance, biological considerations such as lineage-specific high repeat content, extreme GC content, and ploidy changes may need genome quality evaluation in a more phylogenetic context. While we agree that standardised benchmarks are good, the importance of such lineage-specific idiosyncracies must be handled with caution. Experts familiar with the unique biology of the species under consideration will be an essential resource in appropriately evaluating the genomes generated using standardised pipelines.

Another benefit of the data analysis undertaken in this review is careful consideration of the meta-data submitted to the databases. We identify the presence of two BioSamples (SAMEA7522241 and SAMEA7522246) with identical meta information. Although the meaning of this data may be obvious to the primary authors, failure to document this information in an easily accessible resource may mean this information is lost forever and will not be available for future efforts to re-analyse these data. Similarly, we note the discrepancy between the manuscript and the NCBI database regarding whether the organelle genome assembled is complete (circular) or linear (incomplete). The consequences of errors in organellar genome annotation are well documented (Prada and Boore 2019).

High-quality chromosome-level genome assemblies have become an indispensable tool for robust evaluation of a range of evolutionary hypotheses. Therefore, a similar rationale should also motivate genome assembly quality evaluation. The importance of the *M. annua* genome in studying sex chromosome evolution, evaluation of evolutionary strata, prevalence of gene loss, and sex-biased gene expression requires the correct assembly of the sex chromosomes (Veltsos et al. 2019). Molecular studies on the TimeTree database (Kumar et al. 2022) support a median split time of 46 MYA between *Ricinus communis* and *Mercurialis annua*. Individual estimates of 27.4 MYA (Yessoufou et al. 2013), 30.3 MYA (Hermant et al. 2012), 35.8 MYA (Dexter and Chave 2016), 55.8 MYA (Zhang et al. 2022), 75.6 MYA (Cervantes et al. 2016), and 79.6 MYA (Nieto-Blázquez et al. 2017) suggest a split time within the range of 27.4 to 79.6 MYA. High-quality genome assemblies of phylogenetically closer species to *M. annua* will be essential for studying chromosome evolution and robust identification of rearrangement events involved in strata formation (Yazdi and Ellegren 2018).

The advances in genome assembly methodology driven by the availability of long-read sequencing technologies have made it possible to generate chromosome-level assemblies that span telomere-to-telomere while assembling all intervening repeats, including centromeres (Nurk et al. 2022). We argue that all high-quality assemblies should be assessed based on how well the centromere and telomere regions are assembled, as these are the harder-to-assemble regions that can be solved with the latest genome assembly methods. While we identify putative centromeres based on the heterogeneity of the local GC content along the chromosome, an explicit assay to validate these regions as centromeres would be essential (Melters et al. 2013). Without additional validation, it is impossible to rule out that the region identified is simply a collapsed assembly region.

The most convincing evaluation of genome assembly quality is the ability to obtain novel biological signatures unique to the species. The limitations of a genome assembly are often revealed while trying to answer specific questions. A genome that may be good enough to obtain gene sequences may not be well suited to investigate chromosome landscapes due to a lack of long-range contiguity. Our inference of the demographic history of the autosome and x chromosome appears to have a biological signal that is not an artefact of the genome quality. Further work investigating the split times among closely related species using the pseudo-diploid analysis may provide greater clarity regarding the meaning of these inferences. Nonetheless, simulations that evaluate the different evolutionary scenarios that may have contributed to the recent history of these species will need to be performed to get a more definitive answer.

## Conclusion

The preliminary analysis of the Annual Mercury genome presented by Christenhusz et al. suggests the assembly has high contiguity and gene completeness compared to the benchmark datasets and the genomes of closely related species. We urge the authors to carefully scrutinise the metadata, especially regarding Biosample information and organelle genome annotation. The ever-increasing standards of genome assemblies have reached a stage in which the assembly of centromere and telomere regions can be considered routine and a qualifier for any high-quality assembly. The identified putative centromere regions suggest that these regions may be assembled in the genome. We reconstruct the demographic history of the autosomes and X separately as a preliminary check of the assembly quality and find a signal potentially indicative of a biological phenomenon. Further evaluation of this signal, using simulations and comparisons with other closely related species, may reveal some aspects of the evolutionary history of the X chromosome.

## Acknowledgements

We thank Wellcome Open Research for this open peer review opportunity, which includes a signed peer review report published as an open-access document, which provides a permanent identifier (DOI) for the review report so it can be added to the ORCID record and cited independently of the article with the option to obtain the input of colleagues or mentoring early career researchers by naming them as co-reviewers.

## Data accessibility

The data associated with the analysis performed as part of this review are provided here: https://github.com/ceglab/Mercurialis_annua_review.git

## Reviewer expertise

Genome assembly and comparative genomics, as suggested by the publication of the *Mesua Ferrea* plant genome (the first representative genome assembly of the family *Calophyllaceae* (Clusioids), also from the order *Malpighiales*) (Patil et al. 2021).

